# Synaptotagmin-1 and complexin inhibit spontaneous vesicle fusion by masking PIP2, not by clamping SNARE assembly

**DOI:** 10.1101/2025.03.24.644869

**Authors:** Houda Yasmine Ali Moussa, Kyung Chul Shin, Yongsoo Park

## Abstract

Vesicle fusion underlies neurotransmitter release, enabling cellular communication. The rapid kinetics of vesicle fusion, tight docking to the plasma membrane, and coordination by multiple SNARE complexes suggest that SNARE complexes are already pre-assembled before fusion. Synaptotagmin-1 (Syt-1) and complexin (CPLX) have been proposed to clamp SNARE assembly and arrest fusion. However, these models are largely based on studies conducted under low ionic strength and structural analyses utilizing SNARE–Syt-1 chimera conjugates. We propose phosphatidylinositol 4,5-bisphosphate (PIP_2_) as a critical lipid catalyst that facilitates fusion through electrostatic dehydration. Here we show that neither the C2AB domain of Syt-1 nor CPLX-2 has clamping or inhibitory effect on SNARE assembly. Instead, the C2AB domain and CPLX-2 inhibit Ca^2+^-independent vesicle fusion by masking PIP_2_. Our data resolve the long-standing question of increased spontaneous neurotransmitter release in Syt-1 and CPLX knockout neurons, emphasizing PIP_2_ as an electrostatic lipid catalyst for fusion.

**One Sentence Summary:** Syt-1 and complexin inhibit spontaneous fusion by masking PIP_2_.

## Introduction

Vesicle fusion is a fundamental process to release neurotransmitters and hormones in neurons and neuroendocrine cells^1^. Different types of vesicles are specialized to store specific neurotransmitters: synaptic vesicles (SVs) predominantly contain classical neurotransmitters, whereas large dense-core vesicles (LDCVs) mainly store amines, neuropeptides, and hormones^2,3^. Vesicle fusion is orchestrated by soluble *N*-ethylmaleimide-sensitive factor attachment protein receptor (SNARE) proteins^4,5^ and synaptotagmin-1 (Syt-1) serving as a Ca^2+^ sensor to mediate Ca²⁺-dependent vesicle fusion^6,7^.

Given the ultrafast timescale of vesicle fusion^8,9^, the tight docking of vesicles at a distance of 1∼5 nm from the plasma membrane^10,11^, and multiple SNARE complexes to coordinate fusion^12,13^, it is likely that the SNARE complex is partially pre-assembled before fusion in a vesicle docking state. While several models have been proposed to explain SNARE clamping mechanisms involving Syt-1 and complexin (CPLX), most models rely on data obtained under low ionic strength conditions and structural studies using SNARE–Syt-1 chimera conjugates, where the C2AB domain is artificially linked to SNAP-25 via a short linker^14,15^, thereby making the mechanisms of SNARE clamping controversial^6^. Furthermore, although SNARE assembly provides the energy required to overcome the fusion energy barrier, the fact that SNARE complexes are already pre-assembled in a tightly docked state raises important questions about how the energy barrier is effectively overcome to initiate vesicle fusion.

We recently proposed a paradigm shift in understanding vesicle fusion mechanisms, highlighting phosphatidylinositol 4,5-bisphosphate (PIP_2_) as a critical lipid catalyst for vesicle fusion^16,17^. PIP_2_ facilitates fusion by lowering the hydration energy barrier when vesicles are tightly docked through partial SNARE assembly^16,17^. Masking of PIP_2_ by the effector domain (ED) of myristoylated alanine-rich C-kinase substrate (MARCKS) arrests vesicle fusion, stabilizing the vesicles in a state of partial SNARE assembly and tight docking^16^. The vesicle fusion arrest can be reversed by Ca²⁺/calmodulin (CaM) and protein kinase C (PKC), which dissociate the MARCKS ED from membranes, thereby unmasking PIP_2_^17^.

Syt-1 interacts with PIP_2_ primarily through the polybasic region^18^. CPLX also binds to PIP_2_-containing membranes, competing with Syt-1 for PIP_2_ binding^19^. Here, we provide evidence that neither the C2AB domain of Syt-1 nor CPLX-2 has clamping or inhibitory effect on SNARE assembly. Notably, no interaction between the SNARE complex and the C2AB domain is observed under physiological ionic conditions. Instead, the C2AB domain and CPLX-2 inhibit Ca^2+^-independent vesicle fusion by masking PIP_2_. Our data address the longstanding question of why knockout of Syt-1 and CPLX leads to an increase in spontaneous neurotransmitter release, highlighting the role of PIP_2_ as a lipid catalyst for fusion through electrostatic dehydration.

## Results

### No interaction between the SNARE complex and Syt-1 in physiological ionic conditions

Interaction between the SNARE complex and Syt-1 is observed only under low ionic strength and is entirely disrupted under physiological ionic conditions, including the presence of Mg^2+^/ATP^20^. Electron paramagnetic resonance (EPR) spectroscopy demonstrates no detectable interaction between the SNARE complex and Syt-1 in the presence of ATP^21^. Similarly, experiments with PIP_2_-containing nanodiscs using FRET confirm that ATP-induced charge shielding significantly reduces this interaction^22^. However, potential unconventional interaction modes may remain undetected with labeling strategies used in EPR and FRET measurement.

To further confirm no interaction between the SNARE complex and Syt-1 under physiological ionic conditions, we performed fluorescence anisotropy measurement. Anisotropy, which monitors the conformational flexibility of labeled proteins, provides a sensitive measure of interactions regardless of diverse binding modes^23^. The stabilized SNARE complex, comprising syntaxin-1A (residues 183–288), SNAP-25A (cysteine-free), and VAMP-2 fragment (residues 49–96), was reconstituted into liposomes (Lip.) composed of 60% PC, 15% PE, and 25% cholesterol; no anionic phospholipids included. Alexa Fluor 488-labeled VAMP-2 (residues 49–96) served as the fluorescent probe (**Fig. 1a**).

**Figure 1.**
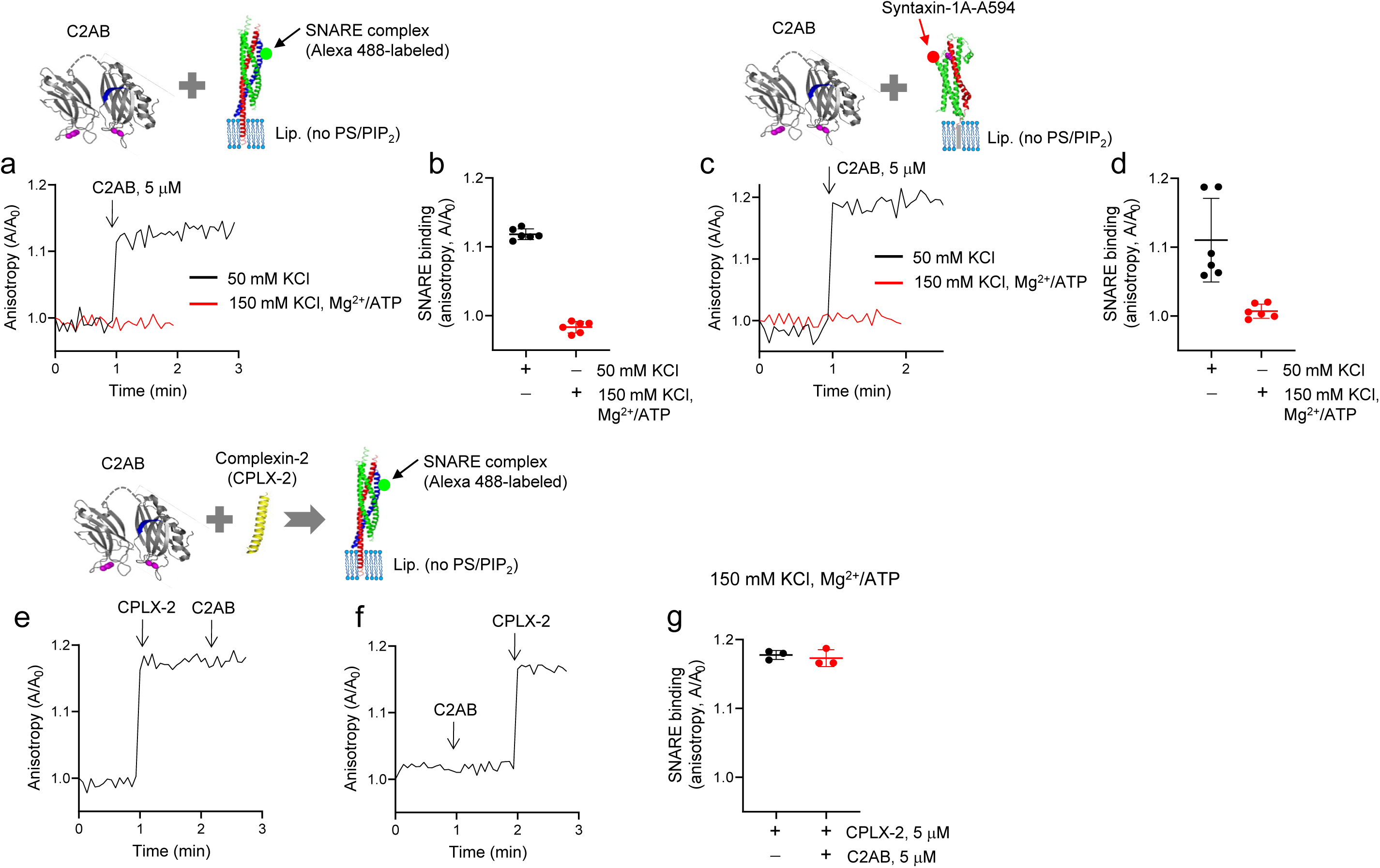
No interaction between the SNARE complex and Syt-1. (**a,b**) Interaction of the SNARE complex with the C2AB domain of Syt-1 was monitored using anisotropy measurement. The stabilized SNARE complex, incorporating VAMP-2 (residues 49–96) labeled with Alexa Fluor 488 (Alexa 488-labeled VAMP-2), was reconstituted into liposomes (Lip.) that contained no PS/PIP_2_: 60% PC, 15% PE, and 25% Chol. The C2AB domain (5 µM) was added as indicated by the arrow. No interaction between the SNARE complex and the C2AB domain was detected under physiological ionic conditions: 1 mM MgCl_2_/3 mM ATP in 150 mM KCl. (**c,d**) Instead of the SNARE complex, full-length syntaxin-1A (residues 1–288) labeled with Alexa Fluor 594 at T197C (magenta) was incorporated into liposomes. (**e-g**) No C2AB domain−SNARE interaction even in the presence of complexin-2 (CPLX-2). Sequential addition of CPLX-2 and the C2AB domain was performed to monitor the interaction with the SNARE complex as described in **a**. Physiological ionic strength with 1 mM MgCl_2_/3 mM ATP in 150 mM KCl was used in **e-g**. Anisotropy was normalized as A/A_0_, where A_0_ is the initial value of anisotropy. Data in **b,d,g** represent means ± SD of three to six independent experiments.

The addition of the C2AB domain of Syt-1 increased anisotropy under low ionic strength conditions (50 mM KCl), confirming SNARE–Syt-1 interaction. However, this interaction was abolished in the presence of 1 mM MgCl_2_/3 mM ATP with 150 mM KCl, demonstrating that physiological ionic conditions disrupt the SNARE–Syt-1 interaction regardless of the different binding modes (**Fig. 1a,b**). Additionally, full-length syntaxin-1A (residues 1–288), labeled with Alexa Fluor 594 at T197C, was tested. The C2AB domain interacted with syntaxin-1A incorporated in liposomes under low ionic strength conditions (50 mM KCl), but this interaction was entirely disrupted under physiological ionic conditions (1 mM MgCl_2_/3 mM ATP with 150 mM KCl)(**Fig. 1c,d**).

Given that CPLX-2 binds the SNARE complex, we further examined the interaction between Syt-1 and the SNARE complex in the presence of CPLX-2. CPLX-2 robustly interacted with the SNARE complex even under physiological ionic conditions (1 mM MgCl_2_/3 mM ATP with 150 mM KCl). However, the C2AB domain showed no additional interaction with the SNARE complex in the presence of CPLX-2 (**Fig. 1e-g**), consistent with previous FRET-based findings^20^. Furthermore, CPLX-2 showed no binding affinity to full-length syntaxin-1A (**Supplementary Fig. 1**). Altogether, we confirmed no interaction between the SNARE complex and Syt-1 in physiological ionic conditions, regardless of the binding modes.

### No clamping effect of Syt-1 and CPLX on SNARE assembly and fusion

To investigate whether Syt-1 or CPLX-2 has a clamping effect on SNARE assembly, we used FRET measurements as previously described^24^. The cytoplasmic domain (CD) of VAMP-2 (residues 1–96) was labeled with Oregon Green as a donor dye, and SNAP-25A was labeled with Texas Red within a stabilized Q-SNARE complex as the acceptor dye (**Fig. 2a**). SNARE assembly was monitored by a decrease in donor fluorescence, which occurs as soluble VAMP-2 CD labeled with Oregon Green assembles with the Q-SNARE complex containing Texas Red-labeled SNAP-25A^25^. The Q-SNARE complex was incorporated in liposomes. As a control, preincubation of unlabeled VAMP-2 CD with the Q-SNARE complex effectively blocked SNARE assembly mediated by labeled VAMP-2 CD (**Fig. 2a**). The C2AB domain did not inhibit SNARE assembly, even under low ionic strength conditions where the C2AB domain interacts with SNAREs (**Fig. 2a, b**). Neither the C2AB domain nor CPLX-2 exhibited any clamping effect on SNARE assembly (**Fig. 2a, b**), despite their interaction with the SNARE complex.

**Figure 2.**
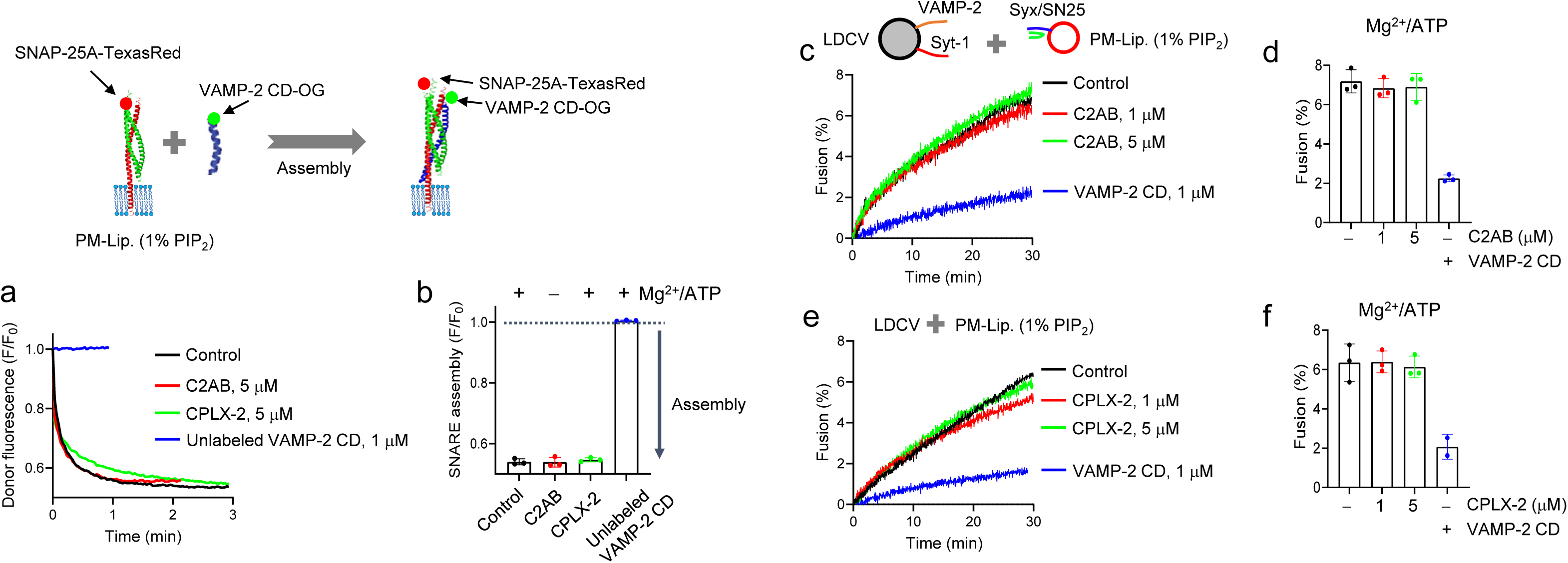
Syt-1 and CPLX-2 show no clamping effect on SNARE assembly and fusion. (**a,b**) The effect of Syt-1 and CPLX-2 on SNARE assembly was investigated using FRET. SNARE assembly was monitored by the cytoplasmic domain (CD) of VAMP-2 (residues 1–96) labeled with Oregon Green and SNAP-25A in a stabilized Q-SNARE complex labeled with Texas Red (refer to **Methods** for details). The VAMP-2 fragment in the SNARE complex was omitted in the diagram for clarity. The reaction was initiated by adding 40 nM labeled soluble VAMP-2 CD to liposomes containing labeled SNAP-25A. SNARE complex assembly led to quenching of donor fluorescence. Liposome lipid composition: 45% PC, 15% PE, 10% PS, 25% Chol, 4% PI, and 1% PIP_2_. Preincubation was performed for 5 min with either 5 µM C2AB domain, 5 µM CPLX-2, or 1 µM unlabeled VAMP-2 CD. Low ionic strength conditions (50 mM KCl, without Mg^2+^/ATP) were used exclusively for the C2AB experiments. FRET was normalized as F/F_0_, where F_0_ represents the initial value of the donor fluorescence intensity. (**c-f**) *In-vitro* reconstitution of vesicle fusion using a lipid-mixing assay in the presence of C2AB (**c,d**) or CPLX-2 (**e,f**). Neither C2AB nor CPLX-2 has the clamping effect on vesicle fusion. Purified native LDCVs were fused with PM-liposomes that incorporated the Q-SNARE complex consisting of the full-length syntaxin-1A (residues 1–288) and SNAP-25A (no cysteine, cysteines are replaced by alanines). The lipid composition of liposomes was consistent with fig. 2a except the addition of labeled PE. Preincubation of PM-liposomes with soluble VAMP-2 CD specifically disrupted SNARE-dependent vesicle fusion. Physiological ionic strength with 1 mM MgCl_2_/3 mM ATP was used in all experiments, unless stated otherwise. Note that ATP removes contaminating Ca^2+^ as a Ca^2+^ chelator^30^ and prevents nonspecific weak interactions between SNAREs and synaptotagmin-1^20^. Data in **b,d,f** are means ± SD from three independent experiments.

Next, we tested whether Syt-1 and CPLX-2 inhibit basal vesicle fusion using purified native vesicles. Physiological vesicle fusion was reconstituted *in vitro* using native vesicles such as LDCVs and SVs under physiological ionic strength (1 mM MgCl_2_/3 mM ATP)^16,17,20,24,26–29^. ATP removes contaminating Ca^2+^ as a Ca^2+^ chelator^30^ and disrupts nonspecific weak interactions between SNAREs and Syt-1^20^, as well as *cis*-interactions of Syt-1 with vesicle membranes^26,30^. Physiological ionic strength with 1 mM MgCl_2_/3 mM ATP is critical for reconstitution of vesicle fusion in a physiological context and therefore, MgCl_2_/ATP were used in all experiments, unless stated otherwise. Native vesicles readily fused with plasma membrane-mimicking liposomes (PM-liposomes, 1% PIP_2_ included) containing the Q-SNARE^31^ (**Fig. 2c-f**). As expected, neither the C2AB domain (**Fig. 2c,d**) nor CPLX-2 (**Fig. 2e,f**) showed the inhibitory effect on basal LDCV fusion, indicating that Syt-1 and CPLX-2 do not clamp or inhibit SNARE assembly and basal vesicle fusion.

### Masking PIP_2_ by the C2AB domain of Syt-1

The data presented above demonstrate that neither Syt-1 nor CPLX-2 shows a clamping effect on SNARE assembly. Given that vesicle fusion can be arrested by masking PIP_2_ in a docking state with partial SNARE assembly^16^, we investigated whether the C2AB domain of Syt-1 inhibits basal fusion by masking PIP_2_. The binding of the C2AB domain to PIP_2_-containing membrane was analyzed using FRET measurement (**Fig. 3**). The C2AB domain (residues 97–421), labeled with Alexa Fluor 488 at residue S342C (donor dye), interacted with liposomes containing Rhodamine-PE (acceptor dye): 45% PC, 13.5% PE, 1.5% Rho-PE, 10% PS, 25% cholesterol, 4% PI, and 1% PIP_2_.

**Figure 3.**
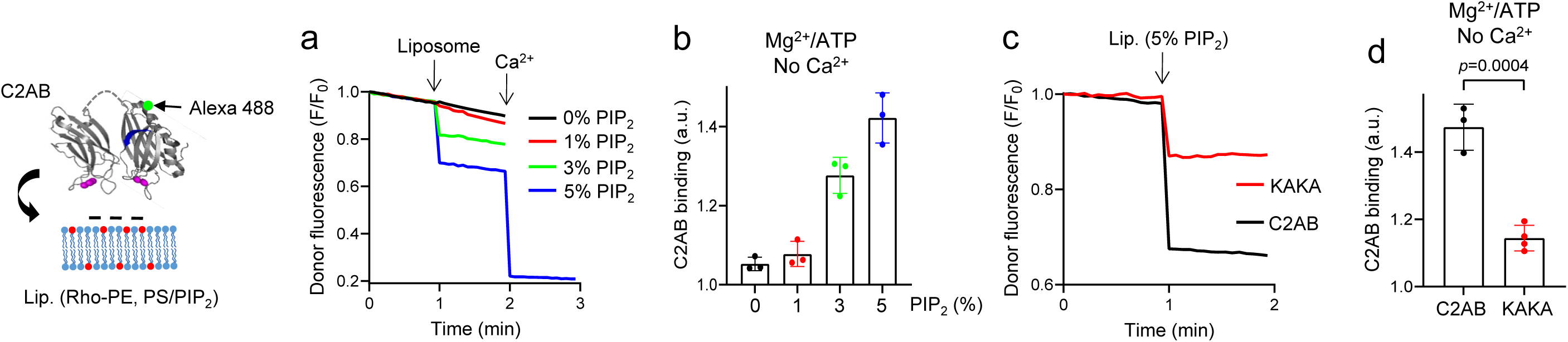
C2AB domain masks PIP_2_ in a dose-dependent manner. (**a-d**) The membrane binding of the C2AB domain was analyzed using FRET measurement. The C2AB domain of Syt-1 (residues 97–421) was labeled with Alexa Fluor 488 at S342C (donor dye, green dot), while liposomes incorporated Rhodamine (Rho)-PE (acceptor dye, red dot). Lipid composition for FRET: protein-free, 45% PC, 13.5% PE, 1.5% Rho-PE, 10% PS, 25% Chol, 4% PI, and 1% PIP_2_. To vary PIP_2_ concentrations, PI content was adjusted accordingly. (**a,b**) Increasing PIP_2_ content up to 5% induced Ca^2+^-independent C2AB membrane binding, whereas 1% PIP_2_ showed little effect. (**c,d**) The KAKA mutant of the C2AB domain reduced Ca^2+^-independent membrane binding. C2AB binding in **b,d** was normalized as F_0_/F, where F_0_ is the initial value of the donor fluorescence intensity. All experiments were conducted under physiological ionic conditions (1 mM MgCl₂ and 3 mM ATP) in the absence of Ca^2+^. Data in **b,d** are means ± SD from three independent experiments. Unpaired two-tailed *t*-test was used (**d**).

The C2AB domain exhibited Ca^2+^-independent binding to 1% PIP_2_ liposomes, but this interaction was disrupted by 1 mM MgCl_2_/3 mM ATP due to charge shielding (**Supplementary Fig. 2**), as reported before^20^. As in a vesicle fusion assay, MgCl_2_/ATP were used to removes contaminating Ca^2+30^ and disrupt nonspecific interactions of the C2AB domain with membranes. Notably, increasing PIP_2_ concentrations to 5% induced robust Ca^2+^-independent binding in a dose-dependent manner (**Fig. 3a,b**). Furthermore, the KAKA mutant of the C2AB domain, in which lysine residues were replaced with alanine (K326A, K327A)^32,33^, showed significantly reduced binding (**Fig. 3c,d**), highlighting the role of the polybasic region in mediating PIP_2_-masking.

### Inhibition of Ca^2+^-independent vesicle fusion by masking PIP_2_

Next, we investigated whether Syt-1 and CPLX-2 inhibit basal LDCV fusion using a lipid-mixing assay. Neither Syt-1 nor CPLX-2 inhibited vesicle fusion when PM-liposomes contained 1% PIP_2_ (**Fig. 2c-f**), consistent with the absence of C2AB binding to membranes containing 1% PIP_2_ in the presence of Mg^2+^/ATP (**Fig. 3a,b**). To evaluate the robust PIP_2_ masking by the C2AB domain, we increased PIP_2_ concentration in PM-liposomes to 5%. Under these conditions, the C2AB domain of Syt-1 inhibited basal LDCV fusion in a dose-dependent manner (**Fig. 4a**). Since CPLX-2 also has PIP_2_-binding affinity^19^, we tested CPLX-2 to reduce basal fusion by masking PIP_2_. As expected, CPLX-2 inhibited Ca^2+^-independent vesicle fusion in a dose-dependent manner, when PM-liposomes contained 5% PIP_2_ (**Fig. 4b**). However, the C2AB domain demonstrated approximately threefold higher potency compared to CPLX-2, with IC_50_ of 0.89 µM for the C2AB domain and 2.99 µM for CPLX-2 (**Fig. 4c**). At the maximum concentration, the C2AB domain and CPLX-2 inhibit basal fusion by ∼50%, whereas MARCKS ED completely blocks basal fusion^16^.

**Figure 4.**
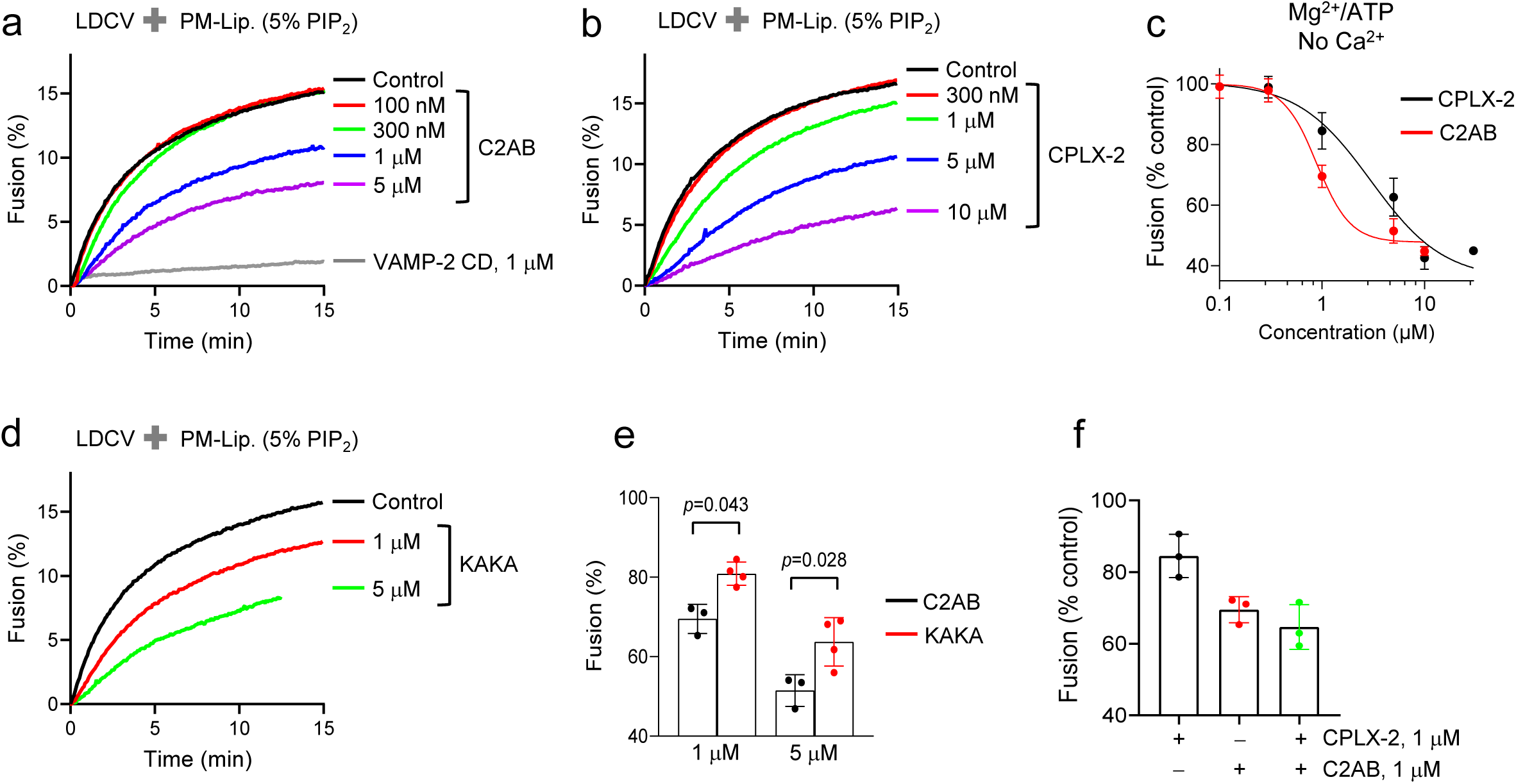
Syt-1 and CPLX-2 inhibit Ca^2+^-independent vesicle fusion by masking PIP_2_. LDCV fusion with PM-liposomes using a lipid-mixing assay. PM-liposomes contained 5% PIP_2_ and incorporated the stabilized Q-SNARE complex consisting of syntaxin-1A and SNAP-25A in a 1:1 molar ratio by the C-terminal VAMP-2 fragment (residues 49–96). The C2AB domain of Syt-1 (**a**) and CPLX-2 (**b**) inhibited Ca^2+^-independent vesicle fusion in a dose-dependent manner. (**c**) The inhibitory effects were quantified, with IC_50_ as 0.89 µM for C2AB and 2.99 µM for CPLX-2. (**d,e**) The KAKA mutant of the C2AB domain showed reduced inhibitory effect on vesicle fusion. (**f**) Co-incubation of 1 µM C2AB domain and 1 µM CPLX-2 did not have additional inhibitory effects on basal fusion. Physiological ionic strength with 1 mM MgCl_2_/3 mM ATP was used. Data in **c,e,f** are means ± SD from three to four independent experiments. One-way ANOVA test with Bonferroni’s post-hoc correction was used.

Further experiments with the KAKA mutant of the C2AB domain, which replaces lysine residues with alanine, revealed reduced inhibition of vesicle fusion compared to the wild-type C2AB domain (**Fig. 4d, e**), supporting the importance of the polybasic region of the C2AB domain in PIP_2_-masking to block basal fusion. Additionally, no significant synergistic effect on basal fusion inhibition was observed when 1 µM of the C2AB domain and 1 µM of CPLX-2 were combined (**Fig. 4f**). Taken together, Syt-1 and CPLX-2 reduce Ca^2+^-independent vesicle fusion by masking PIP_2_, not by clamping SNARE assembly. PIP_2_ is a lipid catalyst to trigger vesicle fusion through electrostatic dehydration, and PIP_2_-masking by Syt-1 and CPLX-2 inhibits basal fusion.

### No effect of Syt-1 and CPLX-2 on SNARE disassembly

Alpha-<u>s</u>oluble <u>N</u>SF <u>a</u>ttachment <u>p</u>rotein (α-SNAP) binds to the Q-SNARE complexes and interferes with full SNARE zippering, thus slowing down fusion and causing tight docking as a result of partially assembled SNARE proteins^24^. The SNARE complex is disassembled by the AAA+ ATPase *N*-ethylmaleimide-sensitive factor (NSF) and a cofactor, alpha-SNAP. Alpha-SNAP first binds to the SNARE complex with a high binding affinity, and then recruits NSF, thereby catalyzing SNARE disassembly in an ATP-dependent manner^34^. In the final set of experiments, we tested whether Syt-1 and CPLX-2 can compete with alpha-SNAP for SNARE disassembly.

SNARE disassembly was monitored using a FRET-based assay, as described earlier^24,35^. Soluble VAMP-2 CD labeled with Oregon Green was assembled with the Q-SNARE complex containing Texas Red-labeled SNAP-25A to form the ternary SNARE complex on liposomes^25^, as in **Fig. 2a**. Alpha-SNAP (500 nM) and NSF (60 nM) were then added in the presence of MgCl_2_ and ATP to activate SNARE disassembly, leading to dissociation of labeled proteins (**Fig. 5a,b**). Disassembly of the SNARE complex was detected by an increase in donor fluorescence intensity. Preincubation with 5 µM of either the C2AB domain of Syt-1 or CPLX-2 did not alter the disassembly of the SNARE complex mediated by alpha-SNAP and NSF (**Fig. 5a,b**). CPLX-2 initially slowed the disassembly kinetics of SNARE complexes, but this effect was rescued within 4 minutes (**Fig. 5a**), because alpha-SNAP binds with higher affinity to membrane-bound SNARE complexes^35^.

**Figure 5.**
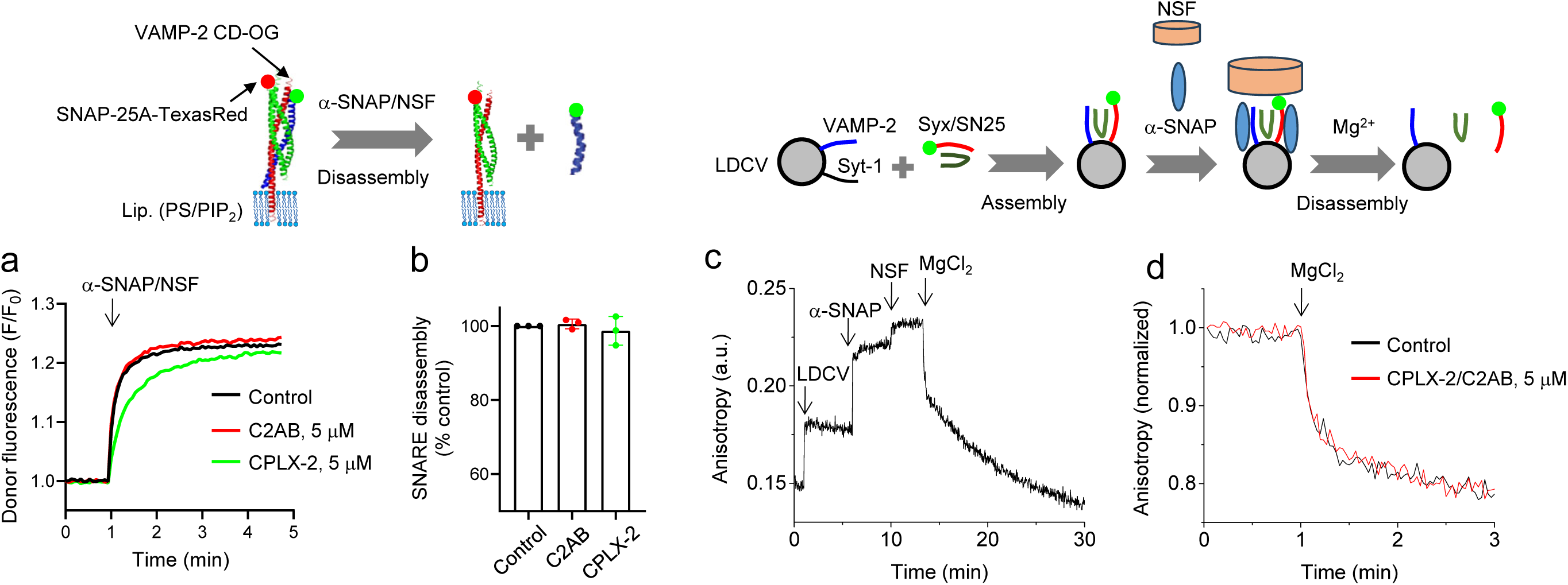
Syt-1 and CPLX-2 do not affect SNARE disassembly. (**a,b**) SNARE disassembly was monitored using FRET measurement. VAMP-2 CD was labeled with Oregon Green and SNAP-25A in the Q-SNARE complex was labeled with Texas Red. The four-helix ternary SNARE complex was formed by SNARE assembly, as described in Fig. 2a. (**a**) SNARE disassembly was measured as an increase in donor fluorescence intensity, corresponding to the disassembly of the SNARE complex. Alpha-SNAP (500 nM) and NSF (60 nM) were added as indicated by the arrow. Preincubation of the SNARE complex with 5 µM C2AB domain or 5 µM CPLX-2 for 5 min showed no inhibition of SNARE disassembly. Liposome lipid composition: 45% PC, 15% PE, 10% PS, 25% Chol, 4% PI, and 1% PIP_2_. (**b**) SNARE disassembly is presented as a percentage of control. 3 mM MgCl_2_ and 3 mM ATP were included in buffer to activate NSF-mediated disassembly. Data are means ± SD from three independent experiments. (**c,d**) Monitoring SNARE disassembly on LDCV membranes using fluorescence anisotropy. LDCVs were incubated with the soluble Q-SNARE complexes (50 nM) comprising syntaxin-1A (residues 183–262), SNAP-25A, and VAMP-2 fragment (residues 49–96), leading to the assembly of membrane-anchored ternary SNARE complexes. Syntaxin-1A (residues 183–262) in the soluble Q-SNARE complex was labeled with Alexa Fluor 488 at position Cys225. Endogenous VAMP-2 from LDCVs replaced VAMP-2 fragment (residues 49–96) to form the ternary SNARE complex. Sequential additions of LDCVs, alpha-SNAP (500 nM), and NSF (60 nM) resulted in stepwise increases in fluorescence anisotropy signals. SNARE disassembly triggered by MgCl_2_ led to dissociation of syntaxin-1A, observed as a decrease in anisotropy signals. 3 mM ATP was included in buffer and as indicated by the arrow, 3 mM MgCl_2_ was applied to activate NSF-mediated SNARE disassembly. (**d**) Co-incubation of 5 µM C2AB domain and 5 µM CPLX-2 did not alter SNARE disassembly.

To validate these findings, we employed an alternative way of monitoring SNARE disassembly, i.e., fluorescence anisotropy. Endogenous VAMP-2 from LDCVs was assembled with the soluble Q-SNARE complexes (50 nM) to form membrane-anchored ternary SNARE complexes on LDCV membranes (**Fig. 5c,d**). Syntaxin-1A (residues 183–262) in the soluble Q-SNARE complex was labeled with Alexa Fluor 488 at position Cys225. SNARE assembly increased fluorescence anisotropy, confirming the formation of the ternary SNARE complex, as reported previously^24^. The SNARE complex formation on LDCV membranes was further validated using SDS-PAGE (**Supplementary Fig. 3**).

Sequential additions of alpha-SNAP (500 nM) and NSF (60 nM) induced additional stepwise increases in fluorescence anisotropy, reflecting protein binding to the ternary SNARE complex (**Fig. 5c**). Activation of NSF-driven disassembly was initiated by adding MgCl_2_, which stabilizes ATP hydrolysis^24^. Co-incubation of CPLX-2 and the C2AB domain had no effect on SNARE disassembly (**Fig. 5d**), indicating that the high affinity of alpha-SNAP for the SNARE complex effectively outcompete Syt-1 and CPLX-2, when the SNARE complex is incorporated in membranes^35^. Altogether, Syt-1 and CPLX-2 do not influence SNARE assembly or disassembly, even under conditions where they interact with SNARE proteins.

## Discussion

We propose a paradigm shift for the molecular mechanisms of vesicle fusion, emphasizing the role of lipids as catalysts in facilitating this process. PIP_2_ and cholesterol are critical plasma membrane lipids required for vesicle fusion (**Fig. 6a**). PIP_2_ acts as a lipid catalyst for vesicle fusion through electrostatic dehydration^16,17^. Cholesterol catalyzes Ca^2+^-dependent vesicle fusion by strengthening Syt-1-induced membrane bending^25,31^. Vesicle docking is initiated by SNARE assembly, which brings vesicle and plasma membranes into close proximity (**Fig. 6b**). Syt-1 and CPLX-2 inhibit Ca^2+^-independent basal fusion by masking PIP_2_, thereby arresting fusion in a docking state. Notably, neither Syt-1 nor CPLX-2 clamps SNARE assembly. PIP_2_-masking arrests fusion due to the hydration energy barriers^16^. PIP_2_ is a lipid catalyst, because PIP_2_ i) attracts cations that neutralize electrostatic repulsion between membranes and ii) repel water molecules, reducing hydration energy and facilitating membrane fusion (**Fig. 6b**). Upon Ca^2+^ influx, Syt-1 is inserted into the plasma membrane, inducing membrane deformation and bending. Cholesterol plays a critical role in lowering the energy barrier for fusion by stabilizing Syt-1-induced membrane deformation (**Fig. 6c**). The final step involves the formation of a fusion pore, enabling the release of neurotransmitters and hormones (**Fig. 6d**). This model highlights the catalytic roles of lipids in driving vesicle fusion and neurotransmitter release.

**Figure 6.**
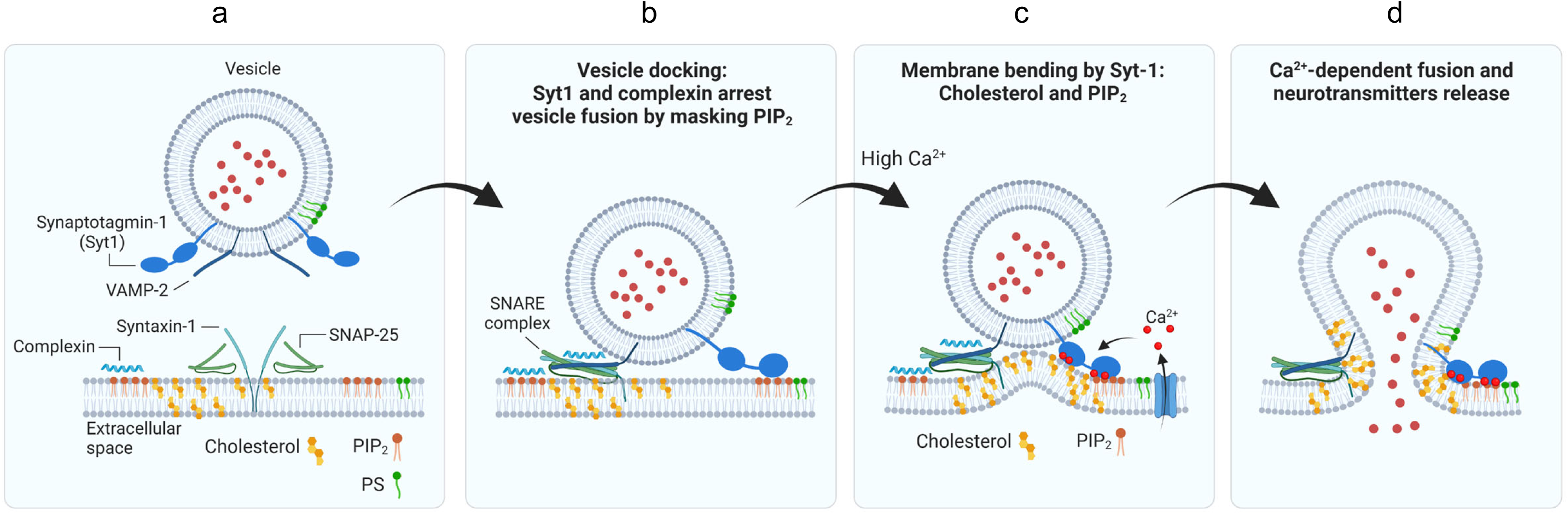
Schematic overview of Syt-1 and CPLX to inhibit Ca^2+^-independent basal fusion. (**a**) PIP_2_ and cholesterol are essential lipids of the plasma membrane for vesicle fusion. PS is present both in vesicle and plasma membrane. (**b**) Vesicle docking is initiated by SNARE assembly, which brings the vesicle and plasma membranes into close proximity. PIP_2_ acts as a lipid catalyst for vesicle fusion by inducing electrostatic dehydration. PIP_2_ attracts cations that i) neutralize the electrostatic repulsion between the membranes and ii) repel water molecules, reducing hydration energy and facilitating membrane merger. Syt-1 masks PIP_2_ and arrest Ca^2+^-independent basal fusion in a state of vesicle docking. CPLX interacts with PIP_2_ on the plasma membrane, similarly inhibiting basal fusion, but without clamping SNARE assembly. (**c**) Syt-1 is inserted to the plasma membrane in response to high Ca^2+^. PIP_2_ induces the *trans*-interaction of synaptotagmin-1 with the plasma membrane. Syt-1 mediates evoked vesicle fusion through membrane deformation and bending, which are strengthened by cholesterol. (**d**) The formation of the fusion pore allows the release of neurotransmitters and hormones.

The SNARE clamping hypothesis was proposed to explain the observed increase in spontaneous neurotransmitter release (mini-frequency) upon deletion of Syt-1 across various cell types^36–42^. Several models suggest that Syt-1 and CPLX act as clamping factors to arrest vesicle fusion, yet these models largely rely on data obtained under low ionic strength conditions, which may not accurately represent physiological environments^6^. Consequently, the mechanisms underlying SNARE clamping remain controversial^6^. For instance, while atomic-resolution X-ray crystallography suggests that Syt-1 interacts with the SNARE complex through a primary interface (reviewed in Ref.^7^), solution NMR studies indicate that the SNARE–Syt-1 interaction is mediated by the polybasic region^43^. These contradictory findings raise further questions about the true nature of the SNARE–Syt-1 interaction. Corroborating earlier reports^20^, we demonstrate that no interaction occurs between Syt-1 and the SNARE complex under physiological ionic conditions, including Mg^2+^/ATP (**Fig. 1a,b**). Using fluorescence anisotropy, we rule out any unconventional SNARE–Syt-1 interactions, regardless of CPLX-2 (**Fig. 1**), arguing against the SNARE clamping model.

EPR spectroscopy provides strong evidence of no detectable interaction between the SNARE complex and Syt-1 in the presence of ATP^21^, consistent with our observations. Similarly, FRET experiments using nanodiscs reveal that ATP-induced charge shielding dramatically reduces SNARE–Syt-1 interaction^22^. The K_d_ of the C2AB domain for PIP_2_-containing nanodiscs is 4.9 ± 1.1 µM, while nanodiscs containing the SNARE complex lower the K_d_ to 0.37 µM, suggesting a robust interaction between the SNARE complex and Syt-1^22^. However, treatment with Mg^2+^/ATP reverses this interaction, increasing the K_d_ to 3.3 µM. Additionally, Mg^2+^/ATP completely disrupts the SNARE–Syt-1 interaction in the presence of 1 mM Ca^2+22^. Our fluorescence anisotropy data further validate no SNARE–Syt-1 interaction under physiological ionic conditions (**Fig. 1**), reinforcing the conclusion that the SNARE–Syt-1 interaction is not physiologically relevant.

We highlighted several factors that lead to experimental artifacts in studies of Syt-1^6^. First, the use of chimera protein complexes in the studies of NMR and X-ray crystallography, where the C2AB domain is artificially linked to SNAP-25 via a short linker^14,15^, can introduce artifacts that mislead data interpretation. To avoid such issues, the soluble C2AB domain should be used to verify SNARE–Syt-1 interactions in structural studies. Second, PIP_2_ should be excluded from membranes, such as nanodiscs, when confirming SNARE–Syt-1 interactions, as the C2AB domain strongly binds to PIP_2_-containing membranes, potentially confounding results. Third, protein structure studies and functional assay should be conducted under physiological ionic conditions, incorporating Mg^2+^/ATP. Syt-1 interactions with membranes or SNARE proteins are tightly regulated by electrostatic effects, making its function and activity highly dependent on ionic strength and salt concentration. Addressing these factors is essential to accurately understand Syt-1 function and interactions under physiological conditions.

Electrophysiology data support our model that Syt-1 inhibits spontaneous vesicle fusion by masking PIP_2_. Substitution of three lysine residues in the polybasic region of the C2B domain with glutamines (K379,380,384Q) increases spontaneous release approximately twofold at the *Drosophila* neuromuscular junction (NMJ)^44^. Similarly, deletion of basic residues (R398,399Q or K326,327E) elevates miniature release frequency and spontaneous fusion in mouse neurons, further supporting the role of PIP_2_-masking in inhibiting basal vesicle fusion^45,46^. Under physiological ionic conditions, the polybasic region of the C2B domain plays a key role in PIP_2_ interaction^18^. Our data support that the KAKA mutation reduces PIP_2_ masking (**Fig. 3c, d**), which correlates with a diminished inhibitory effect on basal fusion (**Fig. 4d, e**). Additionally, R398 and R399 of the C2B domain bind to PIP_2_-containing membranes^43^, and the R398Q,R399Q mutation impairs membrane binding without interfering with the C2AB–SNARE interaction^47^. These findings emphasize the importance of the polybasic region in regulating spontaneous release by masking PIP_2_.

SNARE assembly is essential for vesicle fusion by bringing vesicles into close proximity with the plasma membrane in a docking state, however, our data challenge the hypothesis that SNARE proteins serve as the primary fusion machinery by providing assembly-driven energy to overcome the barriers for vesicle fusion. First, the fact that SNARE complexes are already pre-assembled in a tightly docked state prior to fusion raises critical questions about how the energy barrier is effectively overcome to facilitate membrane merger. Second, Syt-1 and CPLX have no clamping effect on SNARE assembly, and SNARE zippering and vesicle fusion are rarely arrested by SNARE-interacting proteins^16,24^, criticizing the SNARE clamp hypothesis. Third, PIP_2_-masking arrests vesicle fusion in a tightly docked state despite partial SNARE assembly, challenging SNAREs as fusion machinery. Altogether, SNARE assembly is not sufficient to overcome the energy barriers required for fusion. We propose a paradigm shift, identifying PIP_2_ as a lipid catalyst for fusion by lowering the hydration energy barrier.

## Data availability

The datasets that support the findings of this study are openly available.

## Acknowledgements

We thank Dr. Reinhard Jahn for the plasmids and samples. We thank Dr. Ahmed Elalawy and Sarra Karrar from Widam Food Company for the arrangement of adrenal glands. This work was supported by the grant from Qatar Biomedical Research Institute (Project Number SF 2019 004 and IGP5-2022-001 to Y.P.), the HBKU Thematic Research Grant (Project Number VPR-TG02-06 to Y.P.) and the Academic Research Grant (Project Number ARG01-0508-230099 to Y.P.).

## Author Contributions

Conceptualization, Y.P.; Methodology, Software, Validation, Formal analysis, Investigation, Visualization, Y.P., H.Y.A.M., K.C.S.; Supervision, Y.P.; Project administration and Funding acquisition, Y.P.; Original draft preparation, Y.P.; Review and editing, H.Y.A.M., K.C.S.; All authors have read and agreed to the published version of the manuscript.

---

**Supplementary Figure 1.**
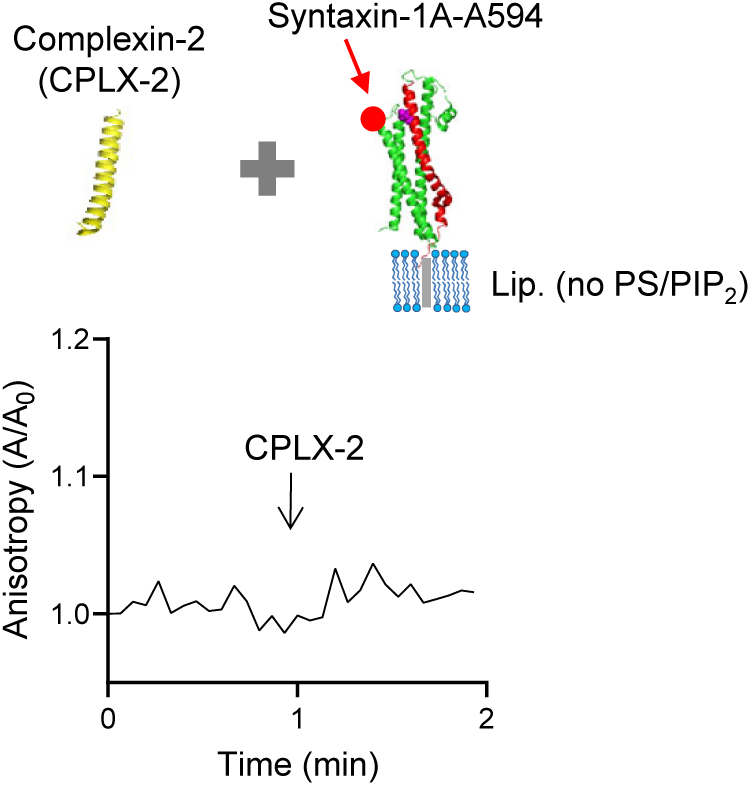
No interaction of CPLX-2 with syntaxin-1A. Interaction between CPLX-2 and syntaxin-1A was monitored using fluorescence anisotropy measurement as described in Fig. 1c. Full-length syntaxin-1A (residues 1–288) labeled with Alexa Fluor 594 at T197C (magenta) was incorporated into liposomes that contained no PS/PIP_2_: 60% PC, 15% PE, and 25% Chol. CPLX-2 (5 µM) was added as indicated by the arrow. No interaction between CPLX-2 and syntaxin-1A was detected. 1 mM MgCl_2_/3 mM ATP in 150 mM KCl. Anisotropy was normalized as A/A_0_, where A_0_ is the initial value of anisotropy.

**Supplementary Figure 2.**
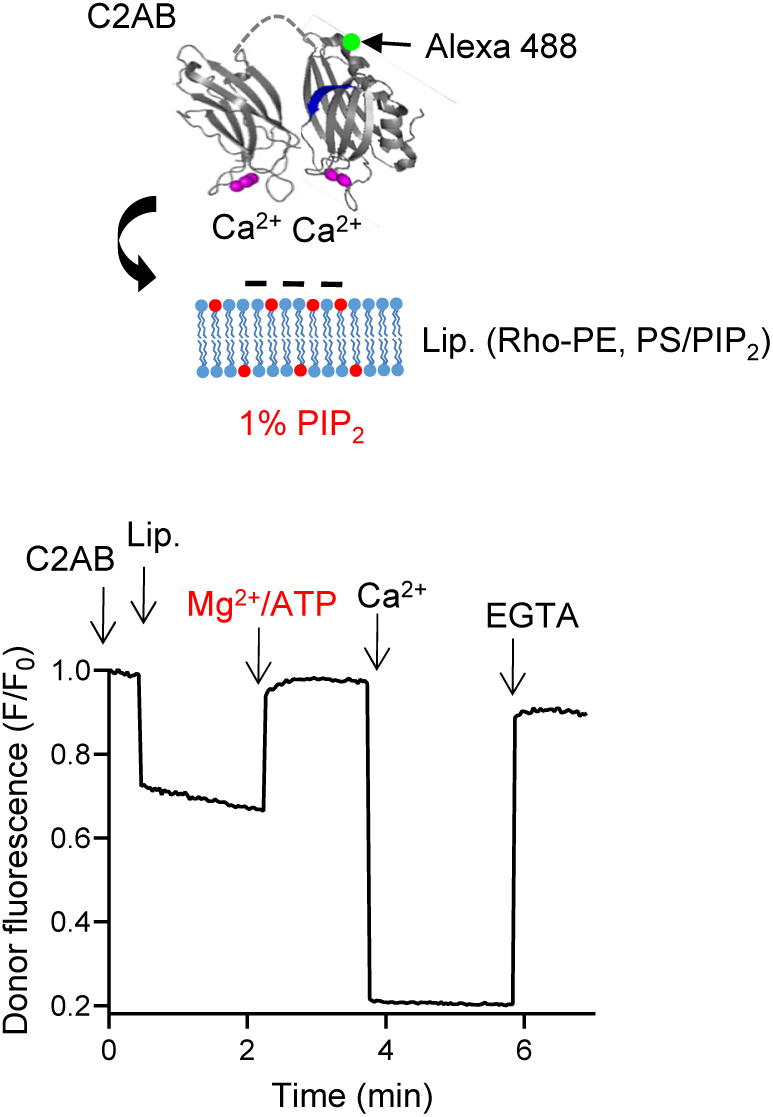
ATP completely disrupts Ca^2+^-independent membrane binding of the C2AB domain. The membrane binding of the C2AB domain was analyzed using FRET measurement as described in Fig. 3. The C2AB domain of Syt-1 (residues 97–421) was labeled with Alexa Fluor 488 at S342C (donor dye, green dot), while liposomes incorporated Rhodamine (Rho)-PE (acceptor dye, red dot). Lipid composition for FRET: protein-free, 45% PC, 13.5% PE, 1.5% Rho-PE, 10% PS, 25% Chol, 4% PI, and 1% PIP_2_. Treatment with 1 mM MgCl_2_ and 3 mM ATP completely disrupted Ca^2+^-independent C2AB binding to liposomes containing 1% PIP_2_. As a control, 100 µM Ca^2+^ induced C2AB membrane binding, which was reversed by 1 mM EGTA. Physiological ionic conditions (1 mM MgCl_2_ and 3 mM ATP) abolished C2AB membrane binding to liposomes that contain 1% PIP_2_. Ca^2+^-independent C2AB membrane binding was observed when membranes contained 3∼5% PIP_2_, as shown in Fig. 3a**,b**.

**Supplementary Figure 3.**
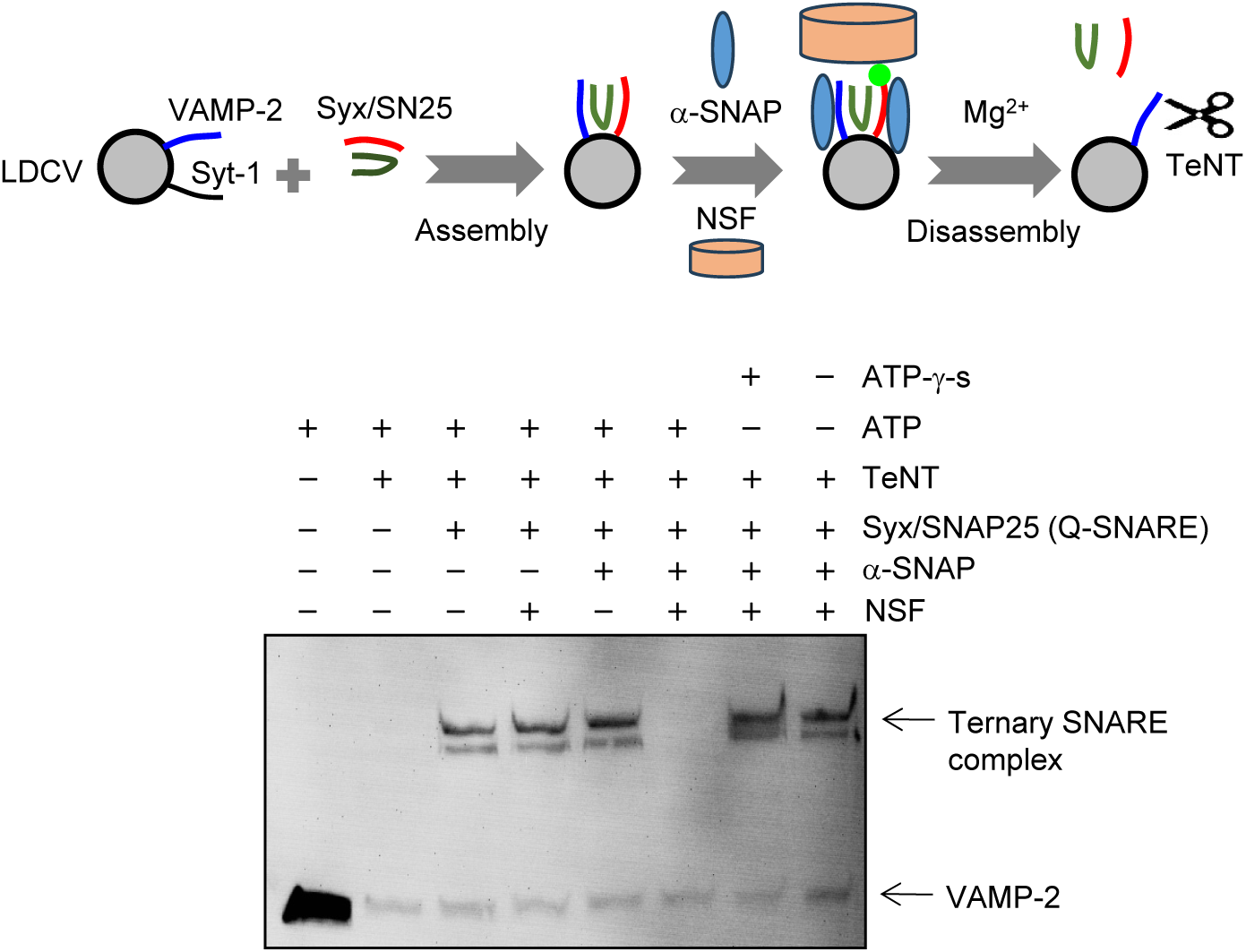
SNARE disassembly mediated by alpha-SNAP and NSF. SNARE disassembly was assessed by analyzing the ternary SNARE complex on SDS-PAGE gels. LDCVs were incubated with the soluble Q-SNARE complexes (50 nM) comprising syntaxin-1A (residues 183–262), SNAP-25A, and VAMP-2 (residues 49–96), resulting in the formation of membrane-anchored ternary SNARE complexes on LDCVs. Endogenous VAMP-2 from LDCVs participated in the formation of ternary SNARE complexes. The light chain of tetanus neurotoxin (TeNT), a protease, selectively cleaves free VAMP-2. VAMP-2 incorporated into the SNARE complex was resistant to TeNT cleavage, migrating as a higher molecular weight band on SDS-PAGE gels, confirming SNARE assembly. The addition of alpha-SNAP (500 nM), NSF (60 nM), 3 mM ATP, and 3 mM MgCl_2_ induced disassembly of the ternary SNARE complex. Degradation of endogenous VAMP-2 by TeNT confirmed SNARE disassembly. Samples were not boiled to preserve the SNARE complex.

## Online Methods

### Materials

ATP disodium salt was from Sigma-Aldrich (Cat. A2383). Oregon Green 488 maleimide (Cat. O6034), Texas Red C2-maleimide (Cat. T6008), Alexa Fluor 594 C5 maleimide (Cat. A10256) and Alexa Fluor 488 C5 maleimide (Cat. A10254) were purchased from Thermo Fisher Scientific. All lipids were purchased from Avanti Polar lipids. Antibody against VAMP-2 (Cat. 104 318, clone number 69.1) was from Synaptic Systems (Göttingen, Germany).

### Purification of large dense-core vesicles (LDCVs)

LDCVs, also known as chromaffin granules, were purified from bovine adrenal medullae by using continuous sucrose gradient and resuspended with fusion buffer containing 120 mM K-glutamate, 20 mM K-acetate, and 20 mM HEPES.KOH, pH 7.4, as described previously^28^. Briefly, fresh bovine adrenal glands were obtained from a local slaughterhouse. The cortex and fat were removed, then the medullae were minced with scissors in 300 mM sucrose buffer (300 mM sucrose, 20 mM HEPES, pH 7.4 adjusted with KOH), and homogenized using a homogenizer. After centrifugation at 1,000g for 15 min at 4 °C, the pellet containing nuclei and cell debris (P1) was discarded. The supernatant (S1) was further centrifuged (12,000g, 15 min, 4 °C), then subjected to an additional cycle of resuspension and centrifugation as a washing step. The resulting pellet (P2, crude LDCV fraction) was resuspended in 300 mM sucrose buffer and loaded on top of a continuous sucrose gradient (from 300 mM to 1.9 M) to remove other contaminants including mitochondria. LDCVs were collected from the pellet after centrifugation at 110,000g for 60 min in a Beckman SW 41 Ti rotor, then resuspended with the buffer (120 mM K-glutamate, 20 mM K-acetate, 20 mM HEPES.KOH, pH 7.4).

### Protein purification

All SNARE, Syt-1, and CPLX-2 constructs based on *Rattus norvegicus* sequences were expressed in *E. coli* strain BL21 (DE3) and purified by Ni^2+^-NTA affinity chromatography followed by ion-exchange chromatography as described elsewhere^20,24,26,31,32,48,49^. The stabilized Q-SNARE complex was composed of syntaxin-1A (residues 183–288) and SNAP-25A (no cysteine, cysteines replaced by alanines) in a 1:1 molar ratio by the C-terminal VAMP-2 fragment (residues 49–96) and the soluble Q-SNARE complex with syntaxin-1A (residues 183–262) were purified as described earlier^31^. The soluble cytoplasmic domain (CD) of stabilized Q-SNARE complex with syntaxin-1A (residues 183–262) was purified as described earlier^31^. The binary Q-SNARE complex containing the full-length syntaxin-1A (residues 1–288) and SNAP-25A (no cysteine, cysteines replaced by alanines) was expressed using co-transformation^26^. Soluble cytoplasmic region of VAMP-2 (residues 1–96) and the C2AB fragment of synaptotagmin-1 (residues 97–421), and the KAKA mutant (K326A, K327A) were purified by Mono S column (GE Healthcare, Piscataway, NJ) as described previously^32,49^. The stabilized Q-SNARE complexes were purified by Ni^2+^-NTA affinity chromatography followed by ion-exchange chromatography on a Mono Q column (GE Healthcare, Piscataway, NJ) in the presence of 50 mM n-octyl-β-D-glucoside (OG)^26^. Soluble VAMP-2 lacking the transmembrane domain called VAMP-2 CD (residues 1–96), and C2AB domain of synaptotagmin-1 (residues 97–421) were purified by Mono S column^24^. Chinese hamster NSF and bovine α-SNAP (wild-type) were expressed and purified as described in detail elsewhere^24,35^. CPLX-2 (residues 1–134) was purified as explained previously^48^. Protein structures were visualized with PyMOL; PDB 1BYN for the C2A domain, 1K5W for the C2B domain, 3IPD for the SNARE complex, 3C98 for syntaxin-1A.

### Protein labeling

The point-mutated C2AB domain (S342C)(C2AB-Alexa 488)^32^, KAKA mutant of the C2AB domain^32^, and VAMP-2 (residues 49–96, T79C) in the stabilized Q-SNARE complex^26,31^ were labeled with Alexa Fluor 488 C5 maleimide. Full-length syntaxin-1A (1–288) monomer was labeled with Alexa Fluor 594 C5 maleimide at T197C^20^. The ternary SNARE complex consisting of syntaxin-1A (183– 288), labeled SNAP-25A (Cys130), and VAMP-2 CD (49-96) was purified on Mono Q column (GE Healthcare, Piscataway, NJ). Syntaxin-1A (residues 183–262) in the soluble Q-SNARE complex was labeled with Alexa Fluor 488 C5 maleimide at position Cys225^24^. A single cysteine SNAP-25A mutant (Cys130) in the stabilized Q-SNARE complex was labeled with Texas Red C2-maleimide^20,24^. VAMP-2 CD (Cys28) was labeled with Oregon Green 488 maleimide^24^. For fluorescent labeling, proteins were incubated with 10-fold molar excess of the fluorophores for 2 hr and unbound dye was removed by gel filtration on a PD-10 column. Protein labeling efficiencies were >70∼80% as assessed by UV-visible spectroscopy.

### Lipid composition of liposomes

All lipids were from Avanti Polar lipids, unless stated otherwise. Lipid composition (molar percentages) of PM-liposomes containing the Q-SNARE complex: 45% PC (L-α-phosphatidylcholine, Cat. 840055), 15% PE (L-α-phosphatidylethanolamine, Cat. 840026), 10% PS (L-α-phosphatidylserine, Cat. 840032), 25% Chol (cholesterol, Cat. 700000), 4% PI (L-α-phosphatidylinositol, Cat. 840042), and 1% PIP_2_ (Cat. 840046). In case of removing PS/PIP_2_, PC contents were accordingly adjusted.

For vesicle fusion lipid-mixing assays, 1.5% 1,2-dioleoyl-*sn*-glycero-3-phosphoethanolamine-N-(7-nitrobenz-2-oxa-1,3-diazol-4-yl) (NBD-PE, Cat. 810145) as a donor and 1.5% 1,2-dioleoyl-*sn*-glycero-3-phosphoethanolamine-N-lissamine rhodamine B sulfonyl ammonium salt (Rhodamine-PE, Cat. 810150) as an acceptor dye were incorporated in PM-liposomes (accordingly 12% unlabeled PE).

For FRET measurement using the C2AB domain labeled with Alexa 488, 1.5% Rhodamine-DOPE was included in liposomes as an acceptor dye. Lipid composition of liposomes: protein-free, 45% PC, 13.5% PE, 1.5% Rho-PE, 10% PS, 25% Chol, 4% PI, and 1% PIP_2_.

For FRET measurement using labeled SNARE proteins, liposome lipid composition is as follows: 45% PC, 15% PE, 10% PS, 25% Chol, 4% PI, and 1% PIP_2_.

### Preparation of proteoliposomes

Incorporation of the stabilized Q-SNARE complex into large unilamellar vesicles (LUVs) was achieved by OG-mediated reconstitution, called the direct method, i.e. incorporation of proteins into preformed liposomes^16,20,26,29^. LUVs prepared by the direct method were used, unless stated otherwise. Briefly, lipids dissolved in chloroform were mixed according to lipid composition. The solvent was removed using a dry nitrogen stream in a fume hood to form a lipid film, and then lipids were resuspended in 0.5 mL buffer containing 150 mM KCl and 20 mM HEPES/KOH pH 7.4. After sonication on ice, multilamellar vesicles were extruded using polycarbonate membranes of pore size 100 nm (Avanti Polar lipids) to give uniformly-distributed LUVs with average diameter of 110 nm^29^. After the preformed LUVs had been prepared, SNARE proteins (for PM-liposomes) were incorporated into liposomes by using OG, a mild non-ionic detergent, then OG was removed by dialysis overnight in 1 L buffer containing 150 mM KCl and 20 mM HEPES/KOH pH 7.4 together with 2 g SM-2 adsorbent beads. Protein to lipid ratio in proteoliposomes was 1:500 (n/n).

### Vesicle fusion assay

A FRET-based lipid-mixing assay was performed to monitor native vesicle fusion *in vitro*^20,26–29^. LDCV fusion assays were performed at 37°C in 1 mL fusion buffer containing 120 mM K-glutamate, 20 mM K-acetate, 20 mM HEPES-KOH (pH 7.4), 1 mM MgCl_2_, and 3 mM ATP. ATP should be made freshly before experiments because it is easily destroyed by freezing and thawing. Free Ca^2+^ concentration in the presence of ATP and Mg^2+^ was calibrated using the MaxChelator simulation program. The fluorescence dequenching signal was measured using Fluoromax (Horiba Jobin Yvon) with wavelengths of 460 nm for excitation (Ex) and 538 nm for emission (Em). Fluorescence values were normalized as a percentage of maximum donor fluorescence (total fluorescence) after addition of 0.1% Triton X-100 at the end of experiments.

### Fluorescence anisotropy measurements

Anisotropy measurement^20,24^ was carried out at 37°C in 1 mL buffer containing 120 mM K-glutamate, 20 mM K-acetate, and 20 mM HEPES-KOH (pH 7.4), 1 mM MgCl_2_, and 3 mM ATP, unless stated otherwise. Anisotropy (*r*) was calculated as *r* = (I_VV_ − G × I_VH_)/(I_VV_ + 2 ×G × I_VH_), where I_VV_ denotes the fluorescence intensity with vertically polarized excitation and vertical polarization on the detected emission, and I_VH_ denotes the fluorescence intensity when using a vertical polarizer on the excitation and horizontal polarizer on the emission. G is a grating factor used as a correction for the instrument’s differential transmission of the two orthogonal vector orientations. Liposomes contained no PS/PIP2: 60% PC, 15% PE, and 25% Chol. Anisotropy (a.u.) was presented as A/A_0_, where A_0_ is the initial value. The stabilized Q-SNARE complex (VAMP-2, residues 49–96) and soluble Q-SNARE complex (Syntaxin-1A, residues 183–262) were labeled with Alexa 488; Ex/Em = 488/516 nm. Protein to lipid ratio in proteoliposomes was 1:500 (n/n).

### Fluorescence resonance energy transfer (FRET)

The C2AB domain (40 nM) labeled with Alexa 488 (a donor dye) was incubated with liposomes that include 1.5% Rhodamine-DOPE (an acceptor dye); donor fluorescence signal was measured with wavelengths of 488 nm for excitation and 516 nm for emission^29^. SNAP-25A in the stabilized Q-SNARE complex was labeled with Texas Red at Cys130 (an acceptor dye) and VAMP-2 CD (40 nM) was labeled with Oregon Green 488 at Cys28 (a donor dye). SNARE assembly led to quenching of donor fluorescence with excitation and emission wavelengths for 488 nm and 520 nm, respectively. Donor fluorescence signal was measured at 37°C using Fluoromax (Horiba Jobin Yvon) in 1 mL buffer containing 120 mM K-glutamate, 20 mM K-acetate, 20 mM HEPES-KOH (pH 7.4), 1 mM MgCl_2_, and 3 mM ATP. FRET was normalized as F/F_0_, where F_0_ represents the initial value of the donor fluorescence intensity.

### Determination of tetanus toxin-resistant SNARE complexes

The light chain of TeNT is a protease that selectively degrades free VAMP-2, whereas VAMP-2, partially or fully assembled in the ternary SNARE complex, is resistant to cleavage^24,50^. Incubation of LDCVs with liposomes containing the stabilized Q-SNARE complex for 10 min at 37°C led to the formation of ternary SNARE complex^24^. For the experiments of SNARE disassembly (**Supplementary Fig. 3**), LDCVs were preincubated with the soluble Q-SNARE complex (50 nM) for 30 min at 37 °C to induce the ternary SNARE complex formation with VAMP-2 on the LDCV membrane. The ternary SNARE complex was analyzed by SDS-PAGE and immunoblotting with antibody against VAMP-2. 500 nM α-SNAP and 50 nM NSF were added to induce disassembly of the SNARE complex for 30 min at 37 °C. At this time, 200 nM TeNT should be included together to cleave disassembled free VAMP-2 in order to block SNARE re-assembly. The ternary SNARE complex formation on LDCV membrane caused the shift of VAMP-2 to an SDS-resistant band of higher molecular weight noted the ternary complex, which was resistant to TeNT, whereas TeNT completely cleaved free VAMP-2^51^. The samples were not boiled to observe the ternary SNARE complex.

### Statistical analysis

Data analysis was performed using OriginPro 2019 software (OriginLab Corporation, Northampton, MA, USA) and GraphPad Prism 9 (GraphPad Software, San Diego, CA, USA). Data are means ± standard deviation (SD). One-way ANOVA test with Bonferroni correction was used to determine any statistically significant differences among three or more independent groups. Probabilities *p* < 0.05 were considered significant.

